# NOFSal-MP10: A single hypervariable STR 12-plex for accurate verification of triploidy in Atlantic salmon (*Salmo salar* L.)

**DOI:** 10.1101/2020.12.22.423910

**Authors:** Celeste Jacq

## Abstract

An increasing interest in production of sterile fish in aquaculture requires rapid, accurate and efficient testing for certification of triploidy prior to sale. Further, validation of triploidy can be beneficial, and even required, for accrediting methods of triploid production. A PCR-genotyping approach using a single megaplex of 12 hypervariable STR markers shows accurate and highly repeatable results enabling verification of ploidy in a single test. The NOFSal-MP10 panel contains 12 STR markers mapped to 9 chromosomes with an average of 21.4 alleles per marker for a combined total of 257 alleles based on genotyped samples to-date. The hypervariable nature of these 12 STRs leads to a large probability for three uniquely sized alleles to be observed at each marker, thus providing a rapid confirmation of ploidy based on the count of allele fragments per marker. Further, as a PCR amplification step is involved, this method is robust to DNA quality and quantity, making it suitable for very early determination of ploidy, as early as the eyed-egg stage. Repeat genotyping of positive control diploid Atlantic salmon over two different capillary electrophoretic instruments in different laboratories and with multiple laboratory personnel proves the panel’s robustness to scoring errors with an overall allelic error rate of 0.3% and a false-positive triploid assignment rate of zero. Genotyping of DNA from 1238 eggs and larvae from 18 independent triploid production batches over three years confirmed triploidy in 98% of samples based on a semi-strict criterion of three unique alleles at one or more loci, and 95% based on a strict criterion of three unique alleles at two or more loci.

**Highlights:**

- This paper describes a rapid quantifiable test for validation of triploidy in Atlantic salmon
- The method is highly accurate and repeatable and robust to DNA quality, allowing testing in embryonic stages
- Rapid early testing of triploidy will enable certified sales of triploid eggs for production

## 1. Introduction

### 1.1. Determination of triploidy

An increasing focus on production of sterile fish in Atlantic salmon aquaculture has led to interest in the use of triploids in production (O’Flynn, et al., 1997; Benfey, 2001; Benfey, 2016). This typically requires an extra cost to the sale of eggs and certification of triploidy may be required upon sale of each batch of eggs so the farmers and management regulators can be certain of the ploidy of the product. As such, there is a need for rapid and accurate estimation of triploidy rates from salmon embryos in aquaculture production. Flow cytometry is often used for ploidy estimation in fish (Allen, 1983; Johnson, et al., 1984) as it shows a proportional increase in the nuclear content of cells according to the number of chromosome copies (Small and Benfey, 1987), and this method has been applied to the embryonic stage in salmonids (Lecommandeur, et al., 1994). However, many flow cytometry protocols involve commercial kits and are often expensive on large-scale sampling, further sample viability is a prohibitive issue under suboptimal storage or transport conditions from field sites (Xavier, et al., 2017) and optimal preservation can vary depending on the type of tissue and age of sample (Hubálek and Flajšhans, 2020).

Genetic markers are another option for verification of triploids as they reflect patterns of inheritance, with the polymorphic nature of STR markers being more suitable for this purpose than biallelic markers such as SNPs due to the increased probability for trisomic allele patterns observed in relatively few loci. Recently however, Grashei, et al. (2020) have developed pipelines for accurate triploid calling from 56K axiom array SNP data. However STR genotyping has the potential to be a more rapid approach for routine triploid validation than a SNP array approach, and furthermore is relatively robust to DNA quality and quantity as it utilises PCR, enabling testing from tiny amounts of starting material, even embryonic DNA. STR genotyping is not without its challenges however, particularly if STR markers with few alleles, or with many rare-alleles have a lower probability for amplification of three uniquely size fragments from an individual, and thus false negative ploidy assignment of a true triploid as diploid may occur when the STR panel possesses little allelic diversity. Problems in accurate scoring of alleles can also occur with highly-polymorphic markers. It is common for the smallest amplified fragment at a STR marker to amplify at greater intensity than a larger fragment at the same marker. One reason for this may be due to the speed at which the polymerase extends the amplified fragment, with shorter fragments reaching complete extension faster than larger fragments and thus having a greater chance of becoming a template themselves for the next chain in the PCR. In extreme circumstances this can result in allele drop-out in STR markers with very large differences in the number of repeats (i.e. large allele ranges). Null-alleles also exacerbate the problem of scoring heterozygotes due to failure to amplify a particular allele, resulting in under-estimates of the true heterozygosity at that marker.

For accurate determination of triploids using STR markers, it is optimal that trisomy at a locus is determined by presence of three differently sized fragments and that this is observed over multiple loci to prevent errors in triploidy assignment. Although it may be possible to distinguish triploids from diploids when only two unique fragments are amplified based on the relative intensity of the peaks, this phenomenon, known as “allele dosing” can be difficult to separate from allele drop-out and should be treated with caution as it may result in false-positive results. Glover, et al. (2015) showed that STRs can provide as accurate estimation of triploidy as flow cytometry. However, published STR-based methods used in robust triploidy determination in Atlantic salmon employ amplification of multiple STR multiplex reactions to obtain sufficient allelic diversity to determine ploidy (Ozerov, et al., 2010; Glover, et al., 2015; Jørgensen, et al., 2018; Glover, et al., 2020). This adds costs both in terms of time and labour, and cost of laboratory resources and consumables, and can therefore lead to delays in the sale of egg batches where the sale is contingent on a certificate to verify triploidy. The application of a single hypervariable STR megaplex that can be genotyped using embryonic DNA may solve this issue and provide a rapid and highly efficient test for triploidy validation for the Atlantic salmon aquaculture industry.

### 1.2. Objectives

The objective of this study was to characterise the use and efficiency of a single hypervariable STR megaplex developed for Atlantic salmon as a rapid tool for triploidy determination. The goal was to determine how accurate and consistent the panel can be in confirming triploids, and the upper and lower rates of genotyping scoring success needed to accurately prevent false-positive and false-negative assignment of triploids. The overarching goal was to verify the STR megaplex as a robust method for triploidy validation that will enable high-throughput testing and rapid turn-around times for certification of triploids in aquaculture production.

## 2. Materials and methods

### 2.1. Selection of STR loci and polymerase

A hypervariable megaplex of 12 STR markers “NOFSal-MP10” (Baranski, et al., 2014) was tested for suitability of triploidy in this study. When developing a marker set for use in diverse populations, it is important to be able to accurately identify the allele ranges in samples from a broad geographic range to ensure markers do not overlap. The study of Baranski, et al. (2014) included two megaplexes with 20 novel genome-derived STRs and two previously published STRs. These two megaplexes were broadly screened using DNA from 89 salmon from 19 populations throughout Norway to determine allele boundaries. It is also important to predict the power of the marker set for ploidy validation by observing patterns of heterozygosity and to estimate the frequency of null-alleles. Polymorphic information content (PIC) is the probability that a diploid individual will be heterozygous (i.e. have two different alleles) at a particular marker. In-depth genotyping of the NOFSal-MP10 panel on 2967 wild salmon from 12 rivers in western Norway was performed in relation to other research projects (data not shown) and these genotypes formed the basis of estimates of genetic diversity and polymorphic information content of each marker for this study. The PIC and expected heterozygosity for each of the markers along with the estimated frequency of null-alleles was estimated using the program Cervus (Kalinowski, et al., 2007).

As the goal of this paper was to design a test that can be used for rapid screening, an emphasis was placed on choosing markers that have easy automated scoring capabilities, e.g. that show no, or little stutter patterns, are hypervariable, that show very little bias in allele dosing and very low to no evidence of null-alleles. To minimise the bias inherent in allele dosing and stuttering, Q5^®^ High-Fidelity Hot-Start DNA polymerase (New England Biolabs) was chosen for its fast extension capabilities, very low error rates and blunt-end products (i.e. no A-addition peaks that can reduce automated scoring efficiency) (New England Biolabs, 2020). Electropherograms of the multi-locus genotypes of two diploid parents and their triploid offspring are provided in Figures A.1–A.4 in Appendix A. These figures highlight the morphological characteristics of the marker multiplex and the large peaks with minimal stutter that enable largely-automated scoring procedures.

### 2.2. PCR and Genotyping

DNA extracted from embryonic or larval stages of 18 triploid production batches from 2018-2020 were genotyped for the NOFSal-MP10 array, alongside positive control diploid Atlantic salmon and blank (no DNA) samples. PCR amplifications were done in 5 μL volumes in a 384-format PCR plate using Q5^®^ Hot-Start High-Fidelity 2x Master Mix (New England BioLabs, 2.5 μL per reaction for 1x concentration) and 1 μL template DNA. Relative concentrations of oligonucleotides were optimised to give approximately equal amplification intensity of markers, with a total final concentration of oligonucleotides of 2.51 μM per reaction. Oligonucleotide information including concentration per oligo pair is provided in Table 1, and detailed sequence information including the mapped positions of the markers according to the Atlantic salmon reference genome ICSASG_V2 (Lien, et al., 2016) are provided in Table A.1 in Appendix A. Oligonucleotides in NOFSal-MP10 were designed with relatively high melting temperatures (Tm), thus enabling a 2-step PCR program. PCR cycling conditions are given in Table 2.

**Table 1:**
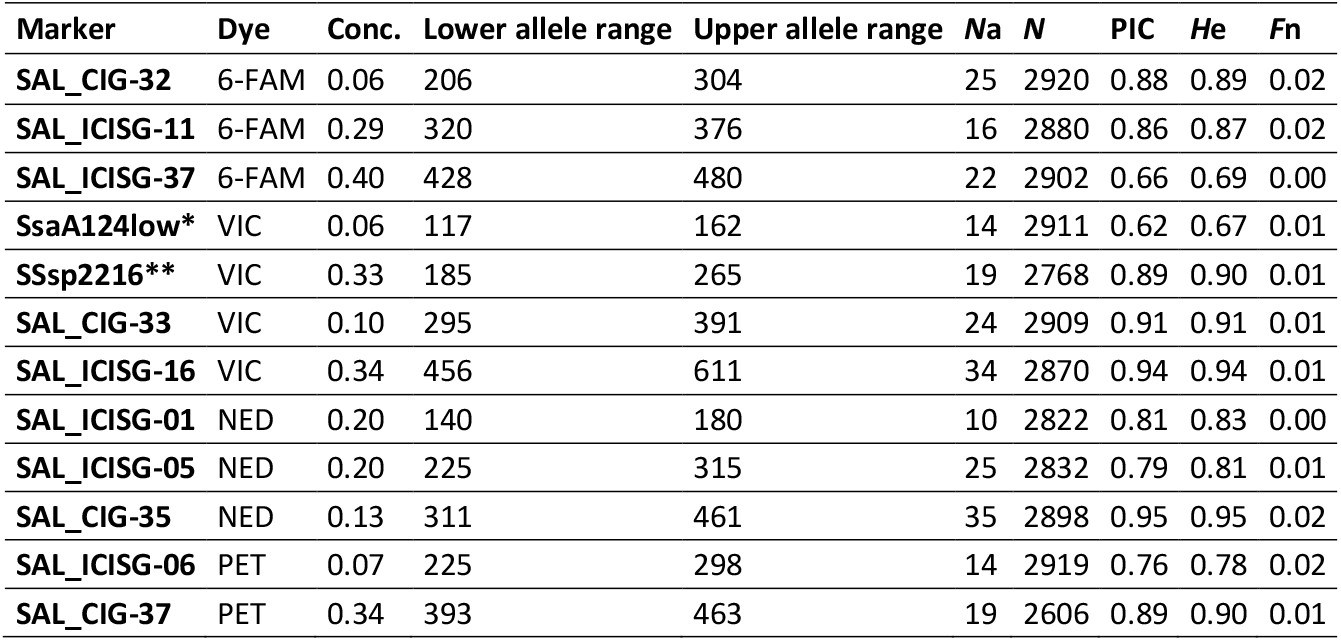
Information for STR markers in NOFSal-MP10. For each marker, the fluorescent dye label used on the forward marker is given (Dye) and the concentration per individual PCR reaction is given in μM (Conc.). The current known extent of the allele ranges, number of alleles (Na), polymorphic information content (PIC), expected heterozygosity (He), and estimated frequency of null alleles (Fn) for Atlantic salmon of European origin is based on population genetic surveys of (N) wild Atlantic salmon from Norway. *redesigned oligonucleotides for SsaA124 (King, et al., 2005); **Sssp2216 oligonucleotides described in (Paterson, et al., 2004).

| Marker | Dye | Conc. | Lower allele range | Upper allele range | Na | N | PIC | He | Fn |
| --- | --- | --- | --- | --- | --- | --- | --- | --- | --- |
| **SAL_CIG-32** | 6-FAM | 0.06 | 206 | 304 | 25 | 2920 | 0.88 | 0.89 | 0.02 |
| **SAL_ICISG-11** | 6-FAM | 0.29 | 320 | 376 | 16 | 2880 | 0.86 | 0.87 | 0.02 |
| **SAL_ICISG-37** | 6-FAM | 0.40 | 428 | 480 | 22 | 2902 | 0.66 | 0.69 | 0.00 |
| **SsaA124low\*** | VIC | 0.06 | 117 | 162 | 14 | 2911 | 0.62 | 0.67 | 0.01 |
| **SSsp2216\*\*** | VIC | 0.33 | 185 | 265 | 19 | 2768 | 0.89 | 0.90 | 0.01 |
| **SAL_CIG-33** | VIC | 0.10 | 295 | 391 | 24 | 2909 | 0.91 | 0.91 | 0.01 |
| **SAL_ICISG-16** | VIC | 0.34 | 456 | 611 | 34 | 2870 | 0.94 | 0.94 | 0.01 |
| **SAL_ICISG-01** | NED | 0.20 | 140 | 180 | 10 | 2822 | 0.81 | 0.83 | 0.00 |
| **SAL_ICISG-05** | NED | 0.20 | 225 | 315 | 25 | 2832 | 0.79 | 0.81 | 0.01 |
| **SAL_CIG-35** | NED | 0.13 | 311 | 461 | 35 | 2898 | 0.95 | 0.95 | 0.02 |
| **SAL_ICISG-06** | PET | 0.07 | 225 | 298 | 14 | 2919 | 0.76 | 0.78 | 0.02 |
| **SAL_CIG-37** | PET | 0.34 | 393 | 463 | 19 | 2606 | 0.89 | 0.90 | 0.01 |

**Table 2:** PCR thermal cycling conditions for NOFSal-MP10

| Step | Temperature | Time |
| --- | --- | --- |
| **Initial denaturation** | 95 °C | 2 min |
| **33 cycles:** | 95 °C | 8 s |
|  | 72 °C | 45 s |
| **Final extension** | 72 °C | 2 min |
| **Hold** | 4 °C | ∞ |

The PCR products were diluted 1:1 with milliQ water and 1μL of the diluted PCR product was added to a 384-format genotyping plate containing 4μL of a mix of Hi-Di™ formamide (Applied Biosystems™) and Genescan™ 600 LIZ™ dye size standard v2.0 (Applied Biosystems™) at a ratio of 96% Hi-Di™ and 4% 600 LIZ™ (v/v). The samples were then analysed via capillary electrophoresis on one of two 3730xL genetic analysers (Applied Biosystems™) with either a 50 cm or 36 cm capillary array.

### 2.3. Genotype scoring and triploidy assignment

Automated allele counts were made using the software Genemarker v.2.6 (SoftGenetics LLC™) with a minimum intensity of 100 RFU required for a peak to be called and where an allele was called for a heterozygous peak that was at least 40% of the height of the other peak at that marker. Manual checks were made for all genotypes to ensure correct allele scoring. Allele counts record the number of uniquely sized allele fragments per marker, and thus take no account of allele dosing or homozygosity.

The allele count table for each batch of genotypes was exported into Microsoft Excel and the genotype scoring rate (GSR) was determined as the number of markers that had an allele count of at least one, divided by the total number of markers genotyped (i.e. 12). Samples were then analysed in respect to triploidy based on the following semi-strict (A), and strict (B) criteria:

A. Where at least one marker had three unique allele fragments.
B. Where at least two markers had three unique allele fragments.

### 2.4. Error rate testing

Two aspects of error testing were undertaken in this study using positive control DNA. For each batch of genotyping, at least two positive control DNA samples were included from a random selection of 13 wild (diploid) salmon with known genotypes. DNA from these fish was genotyped on at least 3 separate occasions, giving a total of 182 genotype profiles with a mean number of 14 multi-locus genotypes per individual. In addition to the GSR which was evaluated in the same manner as for putative triploids, both the false-positive triploid error rate (i.e. occasions that a diploid positive control was falsely assigned as a triploid) and the pairwise allelic error rate (AER) of the STR megaplex were determined from these 182 repeat-genotyped samples. The pairwise AER was calculated from comparisons of the original known genotype and the repeat-genotype of the same individual as: Am/Ac; where Am denotes the number of mismatching alleles and Ac denotes the number of alleles compared. The overall AER for each of the 13 individuals was then obtained as the pairwise AER divided by the number of times the individual was genotyped.

## 3. Results

### 3.1. Triploid validation

DNA from 1600 putative triploids with genotyping scoring rate greater than or equal to 50% was analysed in respect to ploidy with criteria A and B (Figure 1). A greater proportion of triploids were verified using the semi-strict criterion A, and the difference between the proportion of triploids verified between the two criteria increased as the GSR cut-off decreased. The greatest proportion of triploids was achieved at a GSR cut-off 85%-90% for both criteria (97.9% and 94.6% for criteria A and B, respectively), although at a GSR of 80% the difference was marginal (97.7% for criterion A, 94.5% for criterion B) and there was a larger proportion of individuals tested (1238 at GSR 80% compared to 876 at GSR 85-90%). The mean number of markers with three unique fragments for each of these GSR is presented in Figure 2. Although the mean number of markers with three alleles increased as the GSR increased, there was no significant difference between cut-off criteria. Indeed, even with a GSR cut-off of 50%, samples still had on average 4.0 markers with three unique alleles, highlighting the hypervariable nature of this panel of STR markers.

**Figure 1:**
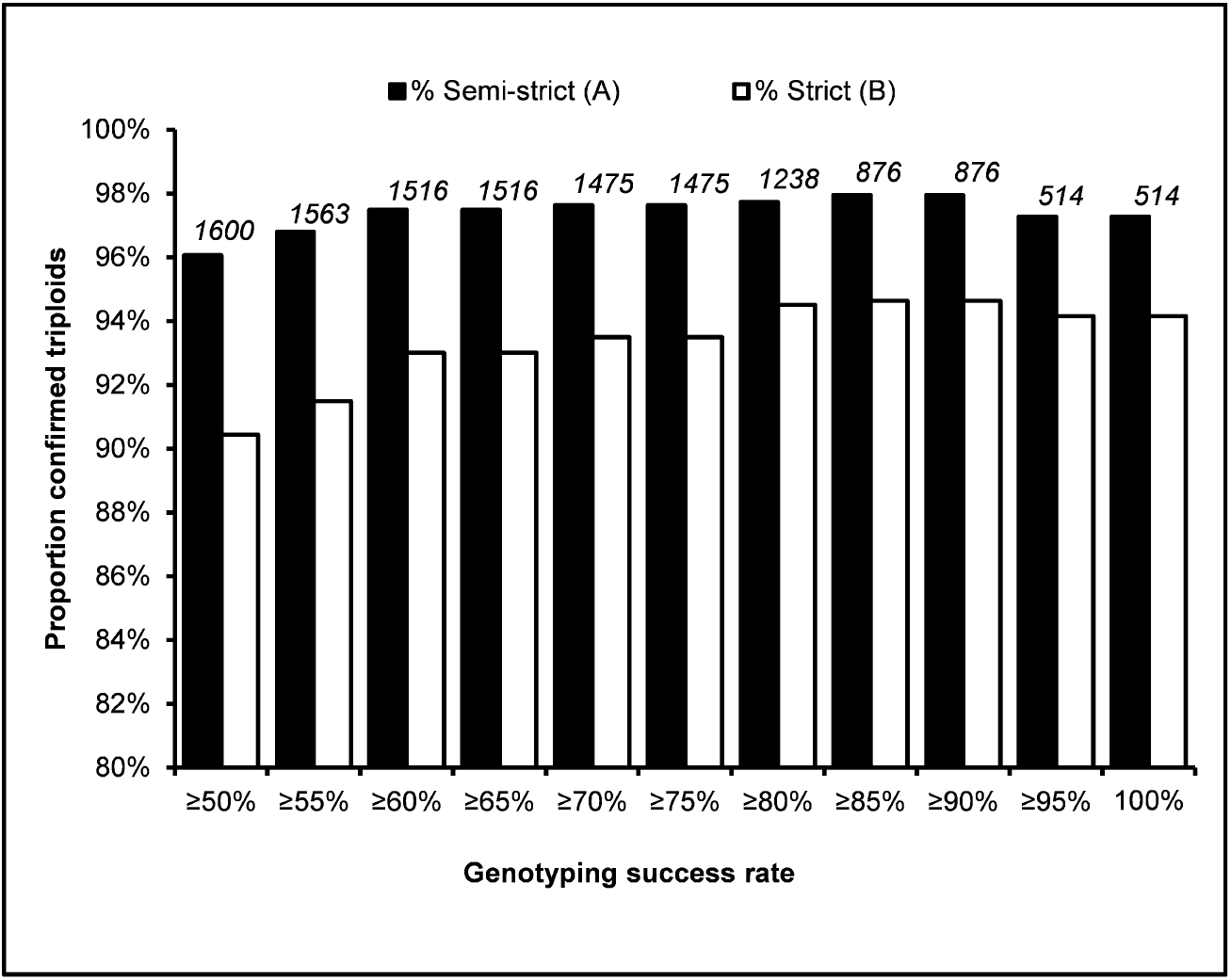
The proportion of confirmed triploid samples evaluated using both semi-strict (A) and strict (B) criteria, given differing cut-off limits to the genotyping success rate (GSR). The number of samples evaluated for each GSR is given above in italics.

**Figure 2:**
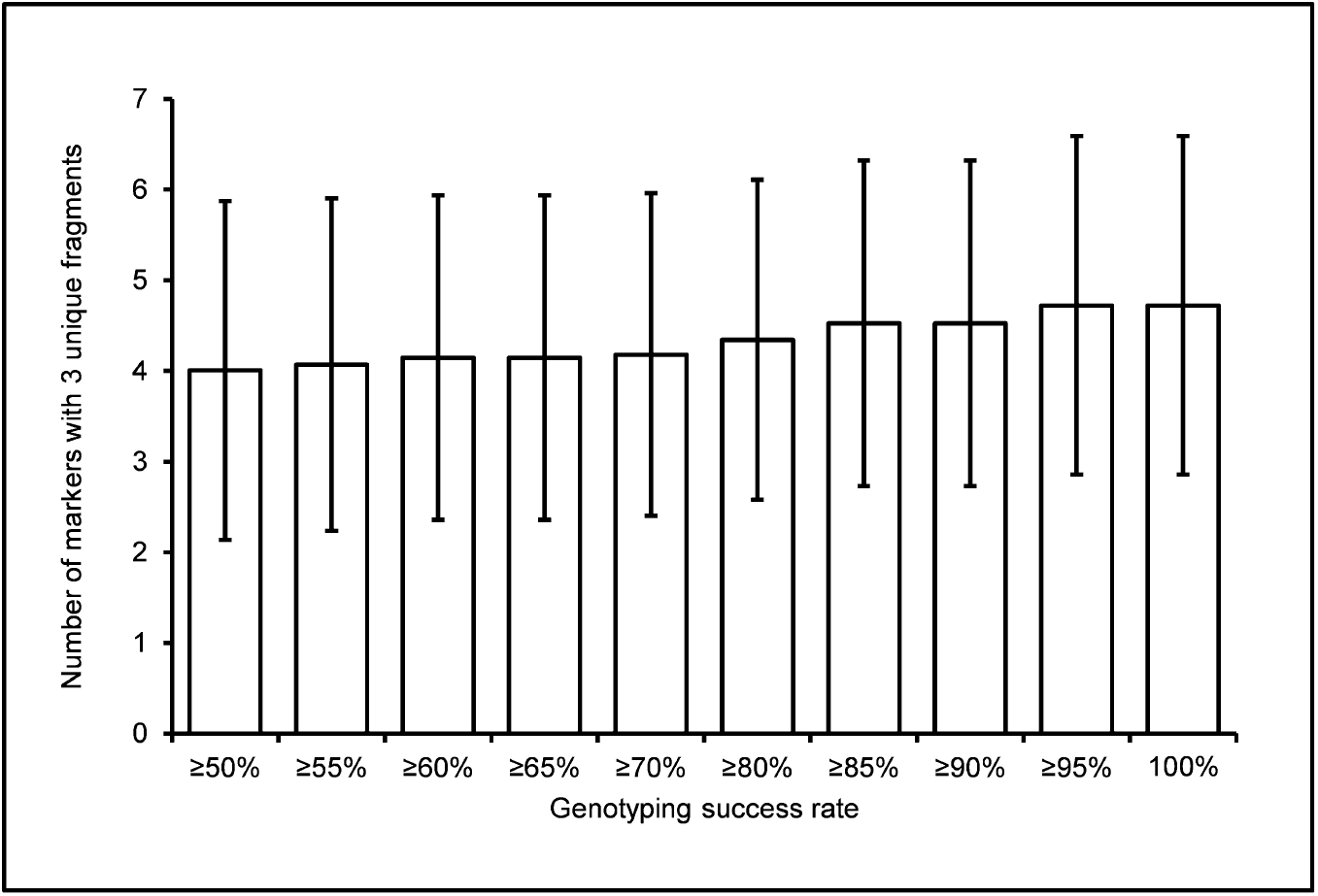
The mean number of markers with three unique allele fragments per individual, given differing limits to genotyping success rate (GSR). Error bars represent the standard deviation

### 3.2. Error rates

No assignment of the diploid control samples as triploids was observed, therefore the false-positive triploid assignment error rate was zero. The average genotyping success rate (GSR) across all positive control genotypes was 98.9% (var. 0.3%, s.d. 5.2%), and the mean calculated multi-locus AER across all 182 repeat-genotyped samples was 0.3%. Allele scoring errors were present in 13 of the 182 multi-locus genotypes and involved pairwise comparisons of DNA from 5 individual fish. No pairwise comparison had more than one allele mismatch across all loci.

## 4. Discussion

Studies on the efficiency and technology involved in producing triploid Atlantic salmon report effective rates of approximately 98% triploids (Benfey, 2016), and this is consistent with triploidy induction rates in other salmonids (Dubé, et al., 1991; Galbreath and Samples, 2000; Devlin, et al., 2010) indicating extremely good, but not perfect effectiveness of the technology. Common methods of triploid induction are through hydrostatic pressure treatment or heat shock soon after fertilisation (Benfey and Sutterlin, 1984), and variation in induction conditions can affect success of the triploid production. The proportion of eggs and larvae from multiple independent triploid production batches verified as triploid in the results presented in this paper are concordant with expected proportions based on published rates of triploid induction in salmonids.

Sub-optimal success of the triploid induction is also known to result in so-called “failed-triploids” and this can manifest in aneuploidy and/or chromosome aberrations (Yamazaki and Goodier, 1993; Devlin, et al., 2010; Glover, et al., 2020). With STR genotyping to validate triploidy, it is therefore vital that the STR markers are positioned on different chromosomes to distinguish possible aneuploids from true triploids. The 12 STR markers in the NOFSal-MP10 panel are mapped to 9 chromosomes on the latest published version (ICSASG_V2) of the Atlantic salmon reference genome (Lien, et al., 2016). Use of the stricter criterion (B), in which more than one marker must have three distinctly sized allele fragments for the individual to be validated as a triploid can be used where distinguishing aneuploids from triploids is required.

If evaluation of triploidy is required prior to sale of eggs, it is essential that DNA from embryos can be tested. Obtaining DNA with high purity and quantity from the embryonic development stages can be difficult, especially earlier in developmental stages. Further, as turnaround time is of importance for the industry, having a test that is robust to errors even with reduced GSR is important. The low GSR inherent in many of these samples is likely due to suboptimal DNA quality and quantity. Automated allele count settings whereby alleles must have an intensity greater than 100 RFU and a heterozygous peak is only scored if it is at least 40% the intensity of the other peak at that locus, ensure strict criteria of peak counting. Results show that the GSR cut-off value affected the proportion of confirmed triploids, particularly with the stricter criteria of three unique fragments at two or more loci, as there were fewer loci included in the evaluation as the GSR decreased. However, lowering the GSR cut-off to 50% had little difference in the proportion of validated triploids using the less-strict criterion A (96.1% at GSR ?50% compared to 97.3% at GSR 100%), even though thrice the number of individuals were included in the evaluation at GSR of 50% or more. This demonstrates the robustness of the validation using criterion A and is due to the semi-strict nature of counting three unique fragments, in addition to the strict allele calling parameters in GeneMarker, especially with low GSR where allele drop-out is more likely. Given these strict calling parameters, it is more likely for an allele to be missed and false negatives to occur, rather than false positives. For validation of triploidy in the aquaculture industry, minimising false positive results is arguably of more importance than minimising false negative results. This is because falsely declaring a diploid individual as a triploid has greater implications both in terms of ethical aspects (selling an item that is not what it actually is), and also environmental effects (diploid individuals are unlikely to be sterile). Therefore, such strict scoring criteria are relevant for minimising false-positive results in assignment of triploids.

Populations that are genetically depauperate typically exhibit greater homozygosity rates in genetic markers. In such populations, validation of triploidy using the methods of three independent fragments at a locus may be difficult due to the reduced probability of an individual having inherited three unique alleles from its parents. Allele dosing refers to the uneven pattern of intensity of alleles when multiple copies of one fragment and a single copy of another are present at the same locus. Triploidy can also be determined by taking account of allele dosing where only two unique allele fragments are amplified per locus, although this method should be treated with caution as a such variation in amplification intensity can also arise due to sequence variation, especially in the priming region of the marker, that may cause less efficient amplification of a particular allele during the PCR. Figures A.1–A.4 show inheritance patterns observed from diploid parents to a triploid offspring. Although seven of the markers clearly show trisomy in the offspring represented as three unique allele fragments, other markers show patterns of trisomy that could potentially be determined by accounting for allele dosing (e.g. SAL_CIG-32, SAL_ICISG-05). However, patterns at e.g. SAL_ICISG-37 could also arise due to allele drop-out in a diploid individual. Thus, triploidy determination by relying solely on allele dosing patterns may lead to false positive assignment of triploids. Nevertheless, the hypervariable nature of the NOFSal-MP10 multiplex, with an overall level of heterozygosity of 0.85 and 257 known alleles to-date, makes this panel a powerful tool for validation of triploidy using an allele count approach rather than an allele dosing approach.

## 5. Conclusion

The NOFSal-MP10 panel described here is an efficient and cost-effective method for validation of triploidy in Atlantic salmon of European origin. The allelic diversity inherent in the marker panel allows relatively strict measures of triploidy validation based on three uniquely sized alleles at one or more loci, negating the need to consider allele dosing to determine triploidy. The robustness of the method to DNA quality allows amplification of DNA from eyed eggs and a genotyping scoring rate of 80% of the markers is sufficient to verify triploidy in proportions consistent to other reported methods, while simultaneously minimising genotyping errors and false positive assignments. The use of a single STR megaplex comprising a fast 2-step PCR approach and small sample volumes ensures an inexpensive test and rapid turnaround time and is also robust to DNA quality and quantity making this a valuable method for validation of ploidy of triploid eggs prior to sale in the aquaculture industry.

## Abbreviations

STR: Short tandem repeat (aka. microsatellite)
GSR: Genotype scoring rate
AER: Allelic error rate

## 6. Acknowledgements

The author would like to thank the relevant industry partners for agreeing to the use of allele count data from genotyped eggs and larvae from triploid production in the analysis. A special thanks go to Katrine Hånes Kirste, Mads Alexander Haneborg, Marianne Helén Selander Hansen, Vibeke Voldvik and Anne Guri Marøy for laboratory assistance.

## Appendix A

**Figure A.1:**
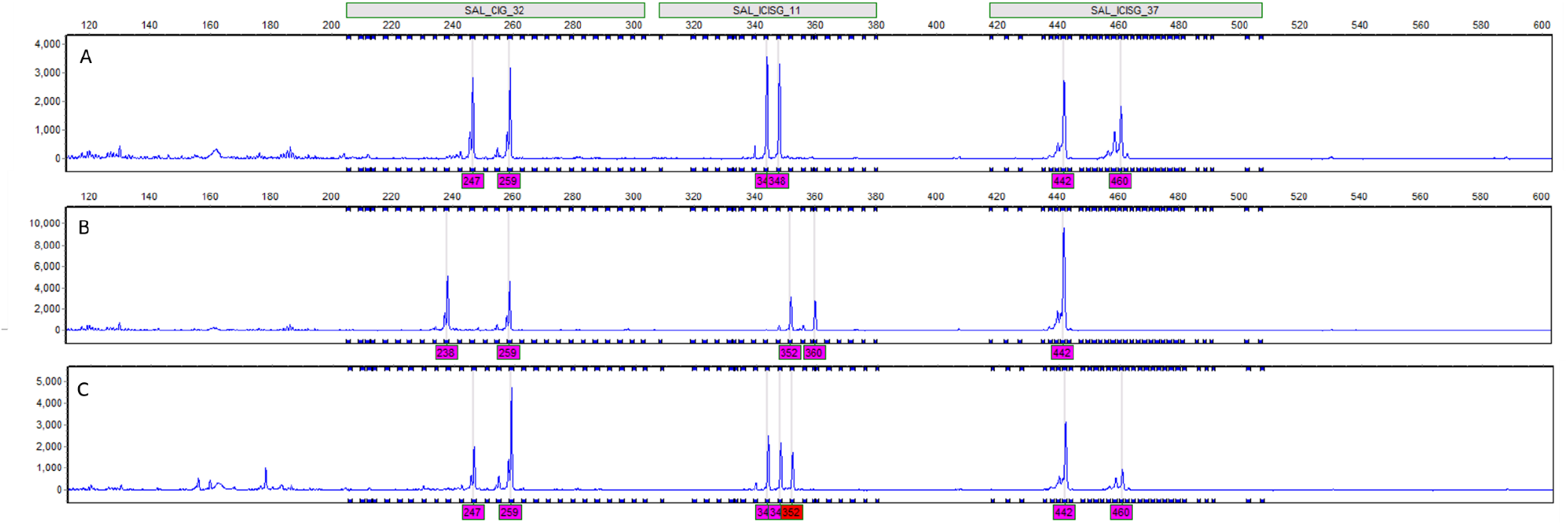
Multi-locus electropherogram output of the 6-FAM dye labelled markers of NOFSal-MP10. Individuals A and B are diploid dam and sire and individual C is their triploid offspring.

**Figure A.2:**
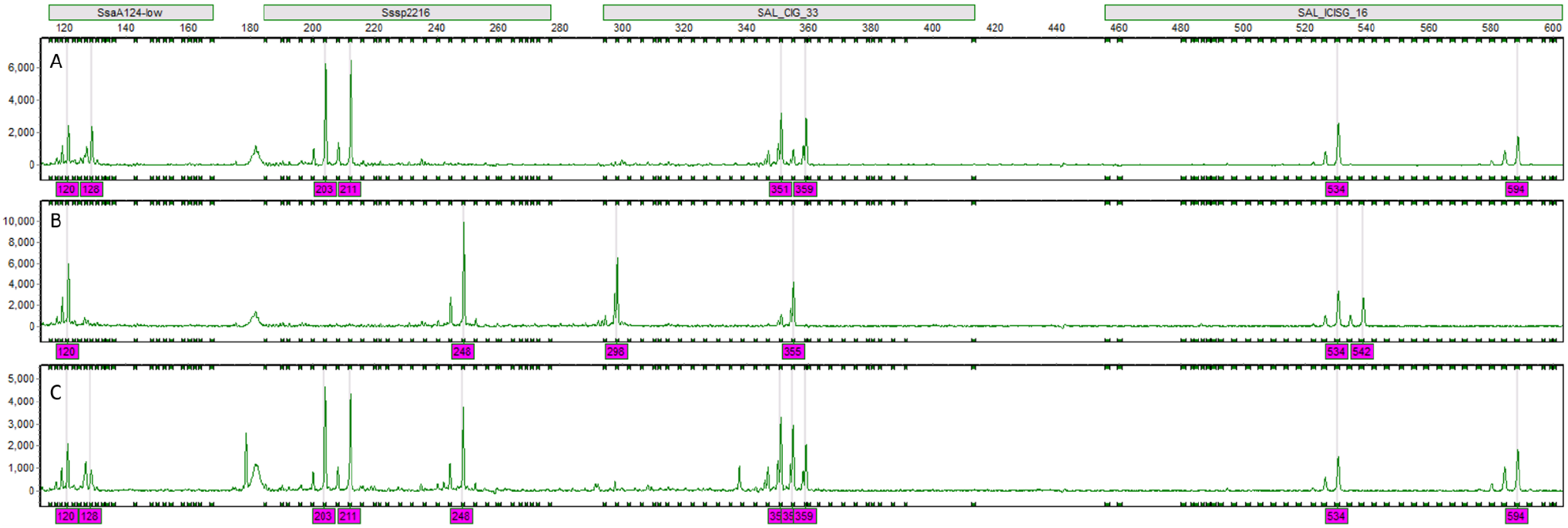
Multi-locus electropherogram output of the VIC dye labelled markers of NOFSal-MP10. Individuals A and B are diploid dam and sire and individual C is their triploid offspring.

**Figure A.3:**
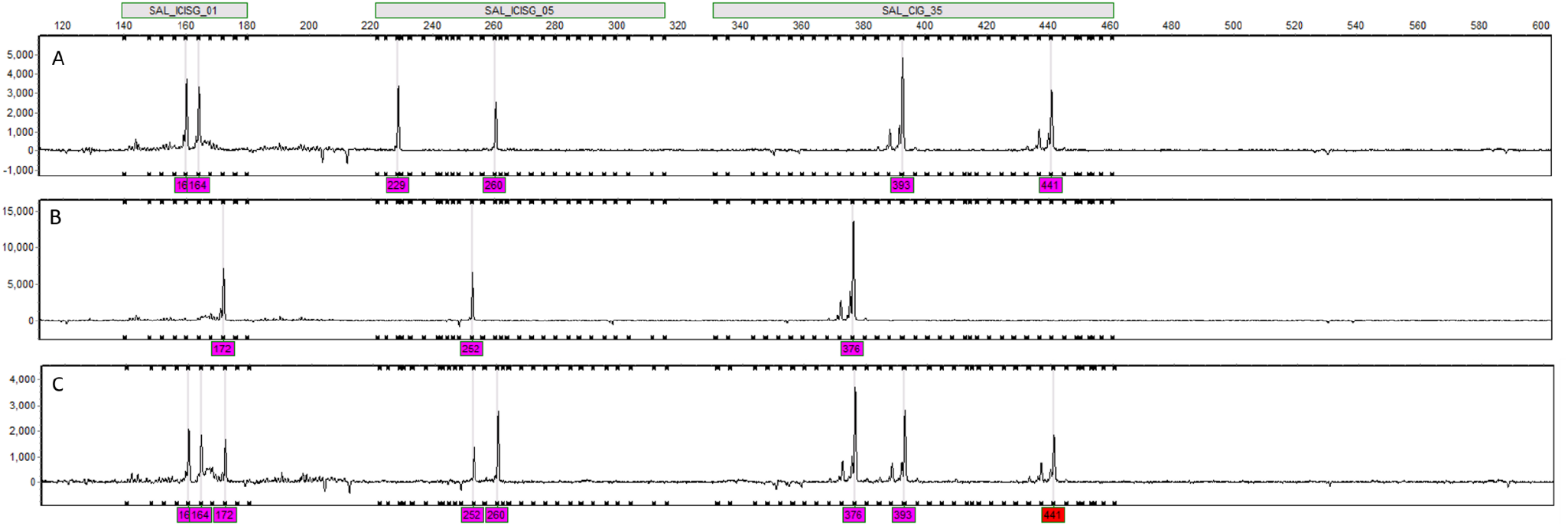
Multi-locus electropherogram output of the NED dye labelled markers of NOFSal-MP10. Individuals A and B are diploid dam and sire and individual C is their triploid offspring.

**Figure A.4:**
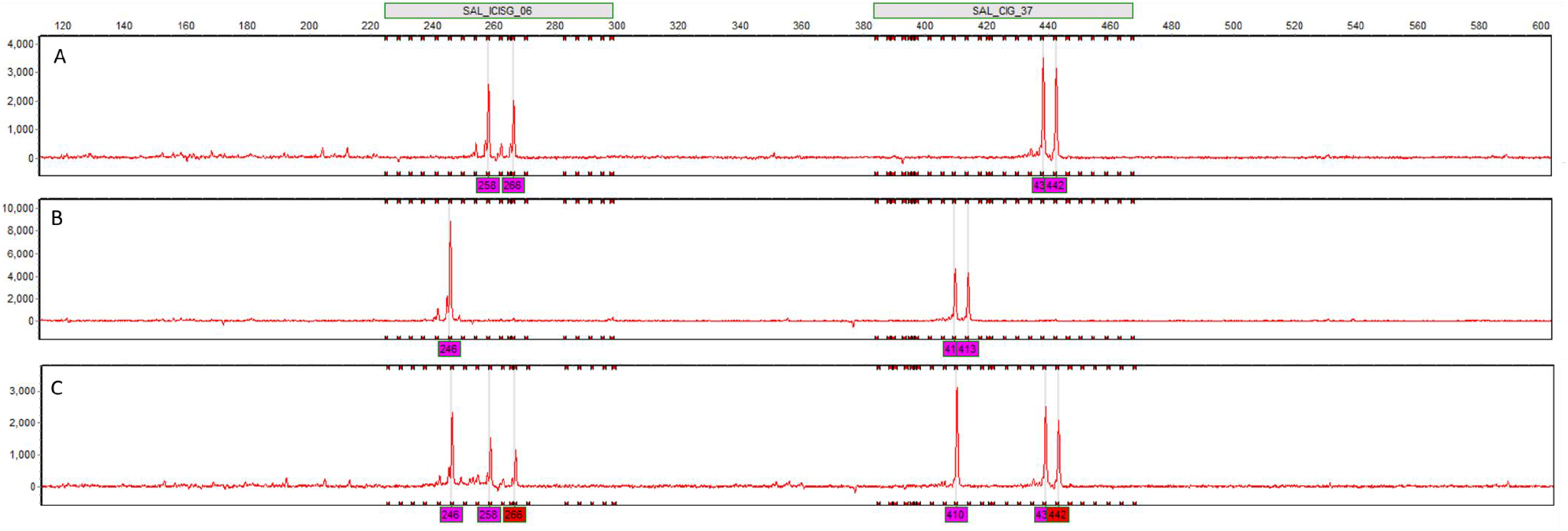
Multi-locus electropherogram output of the PET dye labelled markers of NOFSal-MP10. Individuals A and B are diploid dam and sire and individual C is their triploid offspring.

**Table A.1:**
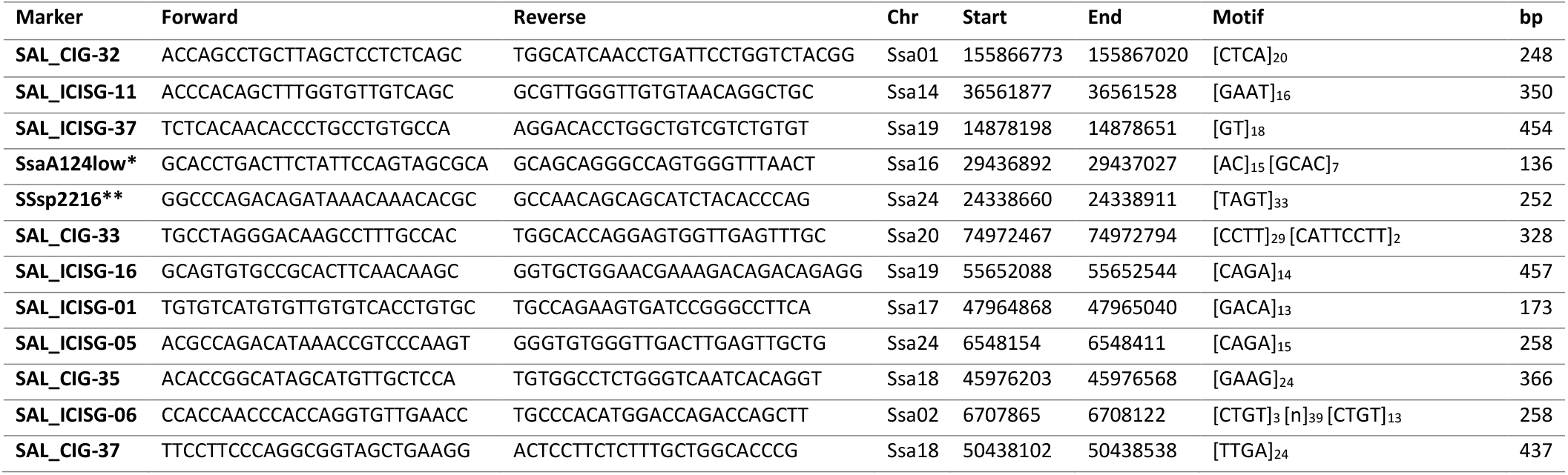
Detailed sequence information for STR markers in NOFSAL-MP10. Forward and reverse oligonucleotide sequences are given along with the coordinates in base pairs (Start, End) on the relevant chromosome (Chr), the repeat motifs (Motif) and amplified product length (bp) according to the Atlantic salmon reference genome (ICSASG_V2, Lien et al 2016). * redesigned oligonucleotides for SsaA124 (King, et al., 2005); ** Sssp2216 oligonucleotides described in Paterson, et al. (2004).

